# Higher Precision in Initial rates may be achievable: A test of a Pseudo-statistical method

**DOI:** 10.1101/2023.04.16.537023

**Authors:** Ikechukwu I. Udema

**Affiliations:** Research Division, Department of Chemistry and Biochemistry, Ude International Concepts LTD (862217) B.B. Agbor, Delta State, Nigeria

**Keywords:** Aspergillus oryzae alpha-amylase (EC. 3.2.1.1), beta-galactosidase (EC.3.2.1.23), maximum velocity, Michaelis-Menten constant, correctional mathematical methods, pseudo-statistical Method

## Abstract

**Background:** There has been a concerted effort at establishing the best method for the measurement of initial rates for various purposes, including the calculation of kinetic parameters, the maximum velocity (*V*_max_), and the Michaelis-Menten constant (*K*_M_).

**Objectives:** The objectives of this research are: 1) to derive equations without *K*_M_ for the determination of the *V*_max_ in particular and *vice versa*; 2) to determine the *K*_M_ and *V*_max_ with other equations other than the Michaelian equation; and 3) to subject the calculated and extrapolated kinetic parameters to pseudo-statistical remediation where necessary as a test of their viability and usefulness.

**Methods:** The study was experimental and theoretical. It is supported by the Bernfeld method of enzyme assay.

**Result:** By graphical means, the *V*_max_ and *K*_M_ values for galactosidase respectively range between 163 and 185 *μ*M/min and between 2.07 and 2.77 mg/L; the range by calculations is 177 and 214 *μ*M/min and 2.45 and 3.311 mg/L, subject to pseudo-statistical remediation. Overall, the ranges of *V*_max_ and *K*_M_ values for alpha-amylase from both the graphical method and calculation are, respectively, 1.095 to 1.018 mM/min and 18.15 to 20.554 g/L.

**Conclusion:** The equations for the determination of the *K*_M_ and *V*_*max*_, which are respectively invariant with respect to each other, were rederived. The initial rates must not be a mixture of both if the true *K*_M_ and *V*_max_ are of interest. The new pseudo-statistical method for the remediation of error in all measurements, if necessary, is viable, useful, and robust.

**Graphical abstract:** A:
Plots where conditions that validate a very high incidence of rQSSA are the case: [*E*_T_] is ≫[*S*_T_].
Plot of *v*_1_ to *v*_5_ versus [*S*_T_]_1_ to [*S*_T_]_5_ gave equation of linear regression (double reciprocal plot (drp)) such as: *y* = (0.08*x* – 0.0002) exp. (+3). A drp plot of all values of *v* versus all values of [*S*_T_] gave a linear regression equation such as: *y* = (0.08 *x* + 6 exp. (− 05)) exp. (+3). 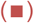 stands for a linear regression of *v* versus [*S*_T_] (*y* = 0.0125*x*); 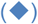 stands for a linear regression of 1/v versus 1/[*S*_T_]. [*S*_T_]_n_*v*_n−1_ − [*S*_T_]_n−1_*v*_n_ is = zero in all data points. The reciprocal of the intercept gives a very high value (over estimation of the maximum velocity, *V*_max_ (16667 mM/min) and consequently an over estimated Michaelis-Menten constant, *K*_M_ (*K*_M_ value is = 106.668 g/L)).

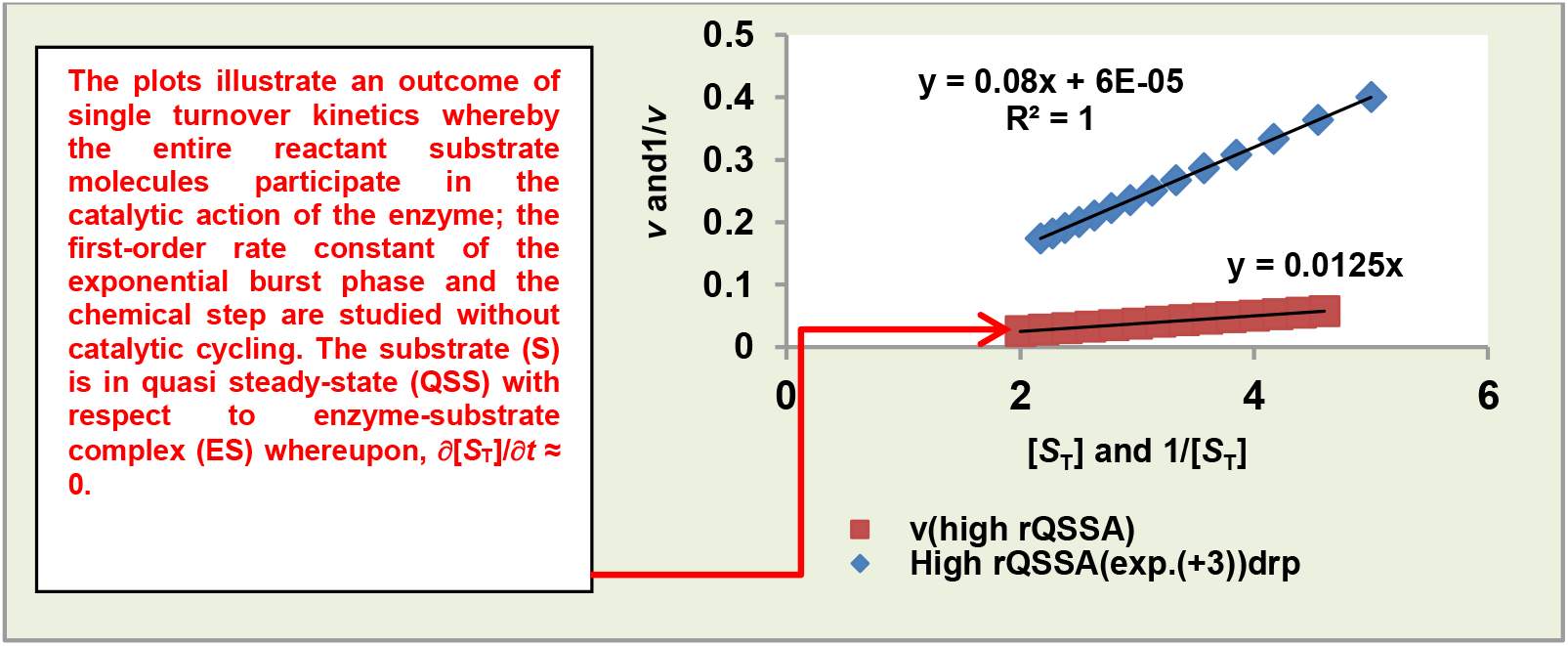

B:
Plots where conditions that neither totally validates an incidence of rQSSA nor sQSSA: Some *v* values are ∝ [*S*_T_] while some are not.
Plot of all *v* values versus all [*S*_T_] values gave equation of linear regression (double reciprocal plot (drp)) such as: *y* = (0.6179 *x* + 0.1973) exp. (+3); the resulting *V*_max_ is = 5.068mM/min and the *K*_M_ is = 3.132g/L. The linear regression of 1/v versus 1/[*S*_T_] gave: *y* = (0.6682*x* – 0.0164) exp. (+3) for the plot covering 1/*v*_1_ to 1/*v*_5_. 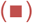 stands for a “polynomial regression” of *v* versus [*S*_T_]; 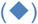 stands for a linear regression of 1/*v* versus 1/[*S*_T_]. [*S*_T_]_n_*v*_n−1_ − [*S*_T_]_n−1_*v*_n_ is ≠ zero where the *v* values covers *v*_7_ to *v*_14_; [*S*_T_]_n_*v*_n−1_ − [*S*_T_]_n−1_*v*_n_ is = zero where the *v* values covers *v*_1_ to v_6_.

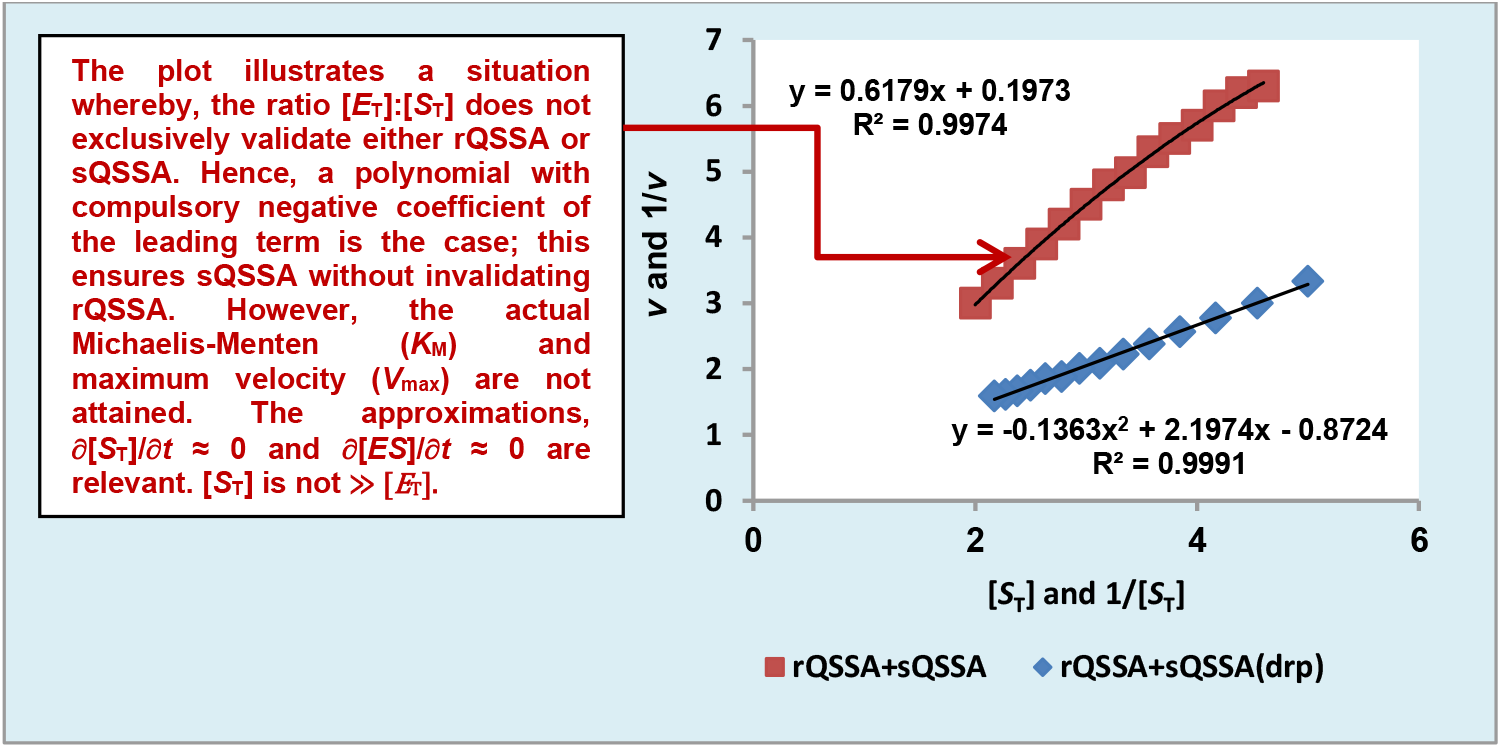

C:
Plots where conditions that validate an incidence of either rQSSA or sQSSA may be the case: Such conditions are [*S*_T_] ≈ [*E*_T_]; [*E*_T_] < *K*_M_.
The *V*_max_ value and *K*_M_ value expected from the regression equation (y = 0.4495 *x* + 0.2921) exp. (+3) from the plot of 1/*v* versus 1/[*S*_T_] are respectively 3.423 mM/min and 1.54 g/L. 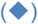 stands for either linear or “polynomial” regression of *v* versus [*S*_T_]: Both plot show *R*^2^ that is = 0.9996; 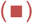 stands for a linear regression (drp) of 1/*v* versus 1/[*S*_T_].

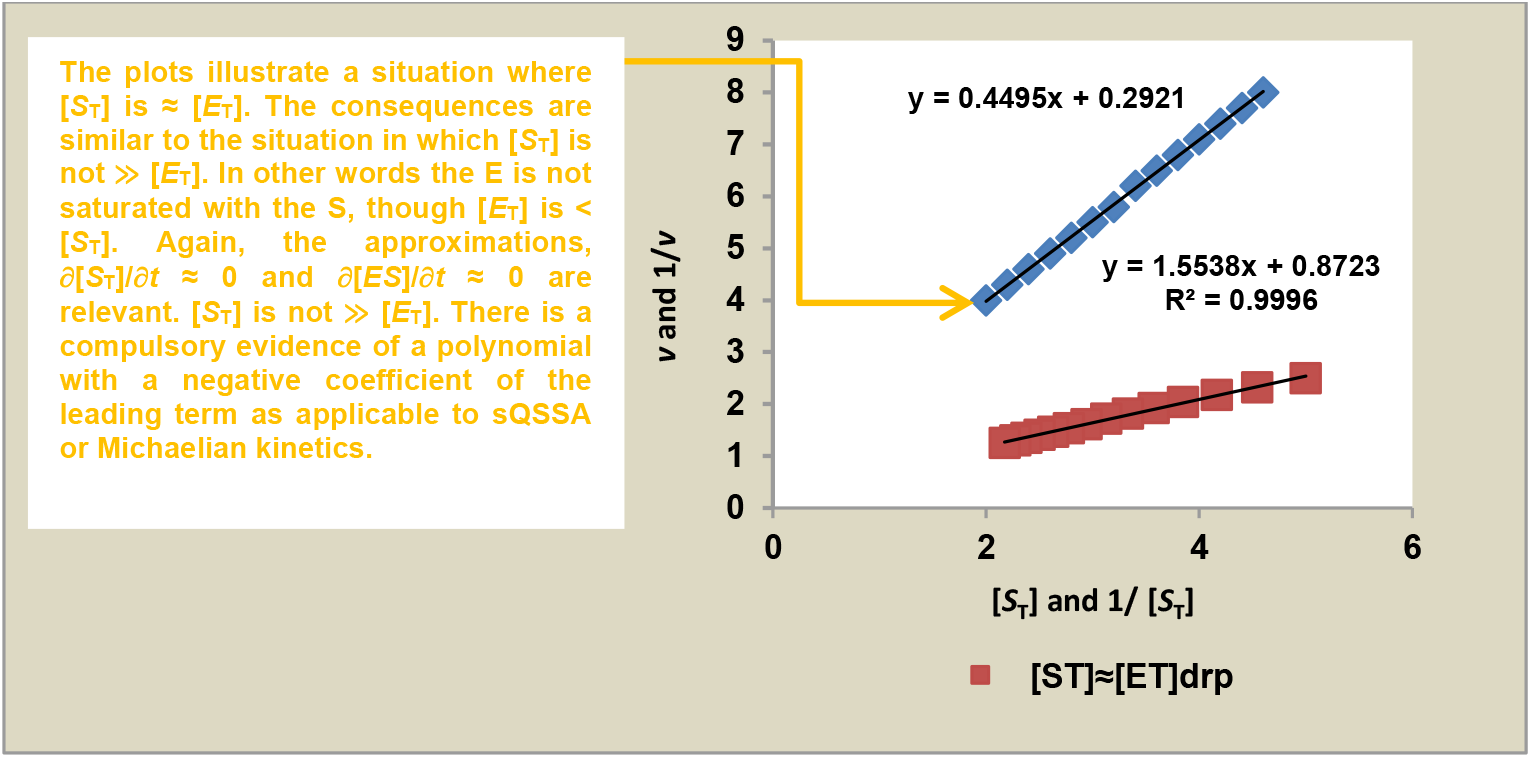

D:
D: Plots where the condition that validate an incidence of sQSSA (or “Henri-Briggs-Haldane-Michaelis-Menten” (HBHMM) equation) may be the case:
Such condition is that [*S*_T_] is ≫ [*E*_T_]. The *V*_max_ value and *K*_M_ value expected from the regression equation (y = 0.0449*x* + 0.0295) exp. (+6) from the plot of 1/*v* versus 1/[*S*_T_] are respectively 33.898 *μ*M/min and 1.522 g/L. 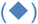 stands for a linear regression of 1/*v* versus 1/[*S*_T_]; 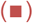 stands for a linear regression of *v* versus [*S*_T_].

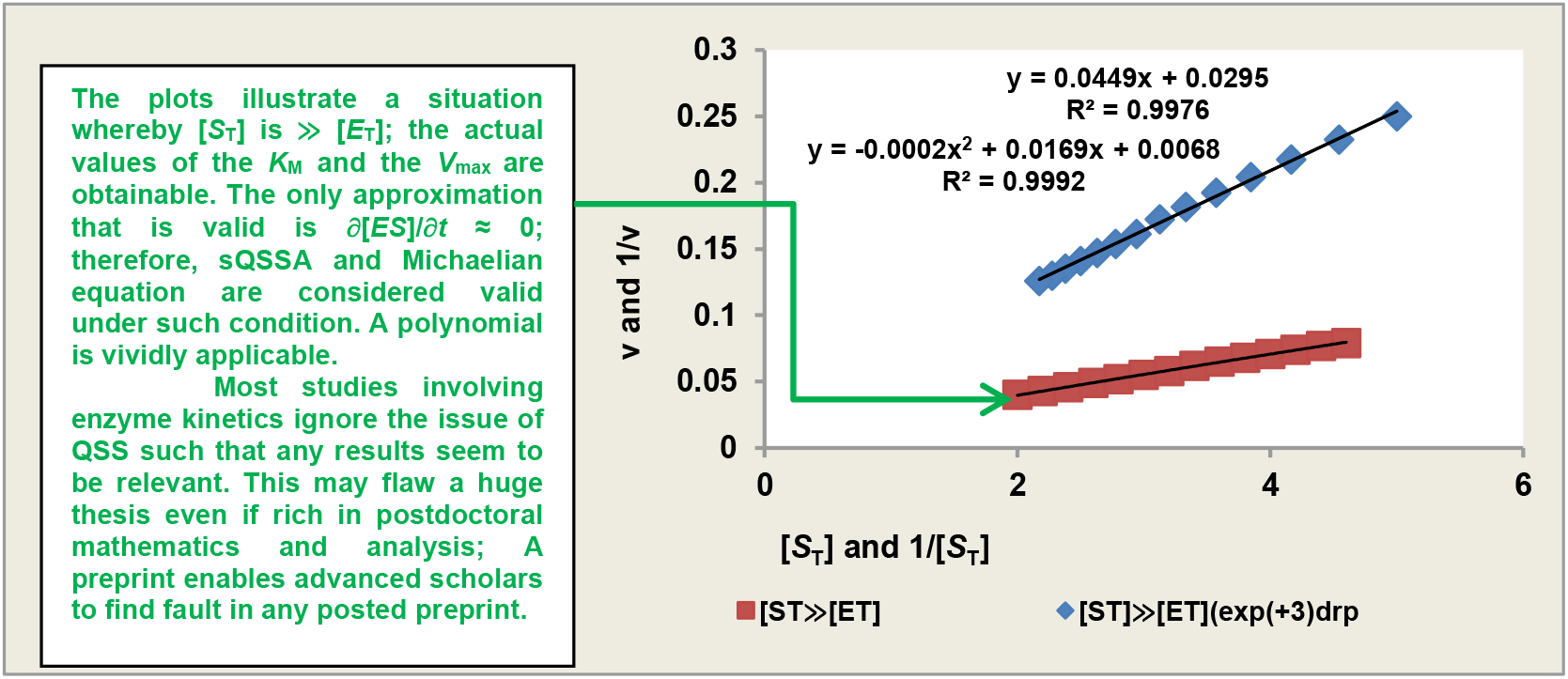

The summary presented in the graphical abstract is primarily intended to remind all and sundry, students and high-ranking scholars in the field, that the issue of QSSA must be reflected in the study of enzyme kinetics because the result of such a study has profound implications for scientific, engineering, and, in particular, medical applications. “To be as imposing as a titanic, does not mean that a titanic-like body is unsinkable”. This implies that minor issues that are ignored can ultimately flaw a post-doctoral thesis by high-ranking researchers. Needless to give an example, but what needs to be taken home is that if an enzyme is very active with a given drug (and even food) to be activated, care should be taken to ensure that a low concentration of drug needs to be administered. In the management of diabetics, starchy foods containing resistant starches are recommended for the same reason.

## 1.0 INTRODUCTION

For more than a century, scientists, the biochemist in the subfield of enzymology, and allied subjects have devoted much attention to the issue of Michaelian kinetic parameter measurement, first executed through the linear transformation of the “Michaelis-Menten [1] equation. However, the latter notwithstanding, Briggs and Haldane [2] played a pivotal role. Also Michaelis-Menten recognised the role of Henri V [3]. To this end, it would have been proper to name the equation the “Henri-Briggs-Haldane- Michaelis-Menten” (HBHMM) equation. A greater motivation for this coinage is reserved for the result and discussion sections. All the while, a hyperbolic curve relating the variation of initial rates with the corresponding concentrations of the substrate has been regularly observed. This implies that the HBHMM model is, *ab initio*, a nonlinear equation, and therefore, it is unwarranted to expect a linear transformation to yield a perfect linear curve even if the data were perfectly generated, *i*.*e*., the total absence of outliers being insinuated. Sometimes, either unknown to the researcher or due to indifference all or two (or more) [4], initial rates may be in a direct proportion to the concentration of the substrate such that the first initial rate (*v*_i_) and its corresponding concentration of the substrate [*S*_T_] are respectively half the next *v*_i_ and the corresponding [*S*_T_]; any double reciprocal plot with the two or more *v*_i_ must create a small negative intercept [5].

According to Matyska and Ková [6], the concerns expressed by enzymologists and statisticians are that the variance σ^2^ of raw experimental data is unknown in most enzymological practice since the experiments are conducted no more than twice, which is not sufficient for the determination of σ^2^. It is therefore necessary to accept some assumptions about the value and structure of this variance in most real experiments. However, assumptions must be treated with strong reservation if applied science, medicine, or safety issues are involved. In a trial-and-error mode, a pseudo-weighting method was developed to bring the raw data much closer to perfection, given a set of rules in place. The concern for the elimination of error has expression in the use of equations such as:

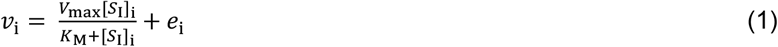

Equation (1) is nothing but the HBHMM equation with an error function, where, as usual, *v*_i_, *V*_max_, *K*_M_, [*S*_T_], and *e*_i_ are the initial reaction rates obtained from steady-state experiments, the maximum reaction rate, the Michaelis-Menten (MM) constant, and random error components. It was not certain how *e*_i_ can be measured.

The best methods of estimating kinetic parameters are, according to Matyska and Kovář [6], the jack-knife Marquardt methods, all of which require a step-by-step approach for adequate comprehension by less gifted scholars in statistics. While *V*_max_ and *K*_M_ can be calculated as intercept and slope from the straight line obtained in a plot of [P]/*t* vs. ln(1- [*P*]/[*S*]*t*)/*t*, the procedure cannot give statistically reliable values of the parameters because the errors associated with [*P*] appear in both the dependent and the independent variable [7]. Thus, most investigations by investigators many years ago [7, 8] were tailored towards the determination of statistical methods for estimating the MM kinetic parameters. The method of least squares gained acceptance with time but gave poor results in the absence of correct weighting, though “bi-weight” regression appeared to be a better option if applied to MM kinetics [9]. This was in response to the failure of almost every linear transformation model to give parameters that are substantially free from errors. This further gave rise to alternative linear transformations: the direct linear transformation model popularised by several researchers [9, 10] and the reciprocal variant [11].

Most, if not all, statistical approaches need statistical packages with which to improve the quality of parameters. If they care, the users of such packages need to be aware of the statistical limitations or validity of **the** weighting routines incorporated into commercially available packages. Perhaps a good example is the R package by Aledo [4]. With the availability of software packages [4, 12], nonlinear regression took centre stage in all attempts to generate reliable Michaelian parameters. Whichever method, a number of substrate concentrations not less than six is required (eight and above is much better) for enzyme assay. This study is therefore, aimed at ways of achieving a higher precision of initial rates given a pseudo-statistical method. On account of the myriad of reservations expressed against various methods, linear transformation in particular, for the estimation of Michaelian kinetic parameters, the objectives of this research are: 1) to derive equations without *K*_M_ for the determination of the *V*_max_ in particular and *vice versa*; 2) to determine the *K*_M_ and *V*_max_ with other equations other than the HBHMM equation; and 3) to subject the calculated and extrapolated kinetic parameters to the pseudo-statistical remediation where necessary as a test of its viability and usefulness.

### 1.1 Significance of study

The subjecting of initial rates to a mathematical analysis in order to identify potential sources of errors that could compromise the quality of the result of the study is very useful; the errors such as direct proportionality between initial rates and the corresponding concentration of substrate leading to negative intercepts in double reciprocal plots suggest an incidence of conditions that justify reverse quasi-steady-state approximation (QSSA), though the condition that validates standard QSSA (sQSSA) is the intention. Further progress demands correctional treatment in line with the methods enunciated in addition to the pseudo-statistical remediation method derived and applied in this research. Fewer replications with concomitant savings in time and material could be an added advantage.

## 2.0 THEORY

Partial reviews of the derivation in a posted pre-print [13] and directly from the usual Michaelis-Menten equation are, respectively:

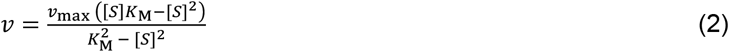

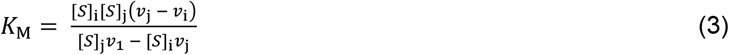

Equations (2) and (3) being general equations lead to the following:

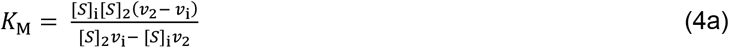

In Eq. (4a), *i* stands for the values of the initial rate and the corresponding concentration of the substrate between the first and the (*n*−1)^th^ sample. The second equation is:

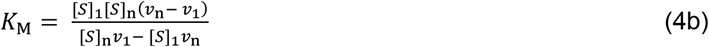

where *i* in the former equation, Eq. (4b), is, in this case, always referring to the first (number 1) initial rate and the first concentration of the substrate, *n* (this could be between 2 and ∞) is always the number of the sample.

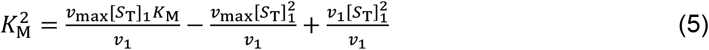

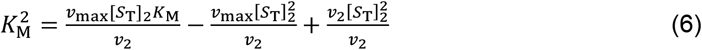

Equations (5) and (6) are the same, so,

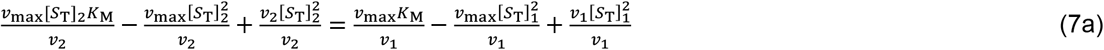

Rearrangement of Eq. (7) gives

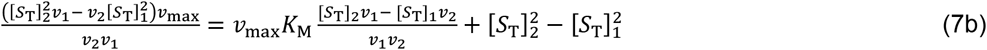

Rearrangement of Eq. (7b) gives:

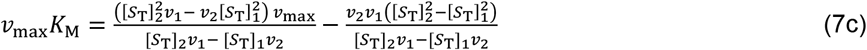

**Equation (4) can now be substituted into Eq. (7c), to give after rearrangement the following:**

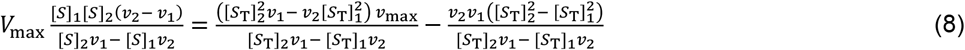

Cancellation of common term or factor and rearrangement gives:

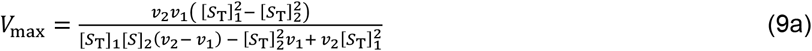

Equation (9a) clearly shows how the *V*_max_ depends on a two-substrate concentration product in both the denominator and the nominator for its calculation. A general equation that should be applied after adjustment in the kinetic variables, following the appropriate equation (s) given in this research, is:

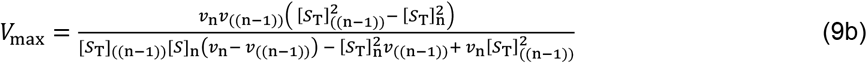

The calculation should cover the variables, *v*_1_, *v*_2_ …, and *v*_n−1_ and the corresponding substrate concentrations, [*S*]_1_, [*S*]_2_ …, and [*S*]_n−1_. The n^th^ variable should be used consistently till the n^th^−1 variable is reached. When the kinetic variables are almost perfectly generated or measured, using high precision equipment, any of the equations for *V*_max_ can be used for its calculation. However preliminary investigation in this research has shown that it is better to adopt Eq. (9a) or equivalent equation in the literature [13] but stated herein shortly because, such enables the earlier disclosure of un-Michaelian trend whereby [*S*_T_]_n_*v*_n-1_ − [*S*_T_]_n−1_*v*_n_ is either negative or zero. Since, in this research, details and a step-by- step approach are matters of policy rather than haste and convenience, another general equation is hereby given as follows:

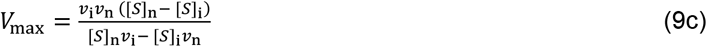

In Eq. (9c) *i* stands for the values of the initial rate and the corresponding concentration of the substrate between the first and the (*n*−1)_th_ sample. Equation (9c) needs to be used to check the first three data. The second equation can take the form:

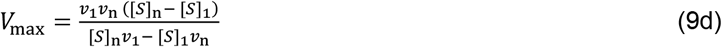

where *i* in the former equation, Eq. (9c) is, in this case, always refers to the first initial rate and the first concentration of the substrate; *n* (this could be between 2 and ∞) is always the number of sample.

The corresponding equation of *K*_M_ is derived as follows. Given the equation in the literature [13], written as below, one can derive the corresponding equation of *K*_M_ as follows:

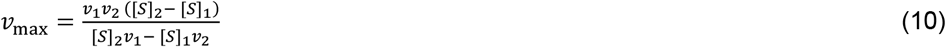

Equation (10) can be substituted into Eq. (7c) to give:

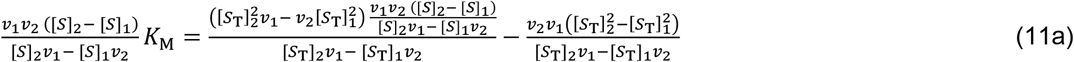

Cancellation of common factors and rearrangement gives:

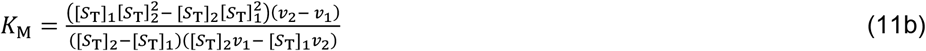

Equation gives exactly the same results when fitted to kinetic variables and substrate concentrations as it is in earlier derivation in the literature [13]. Here the approach partially evaded the direct use of Michaelis-Menten equation, but reaffirmed the procedural validity now and in the past [13]. Again, the general form of Eq. (11b) is given as:

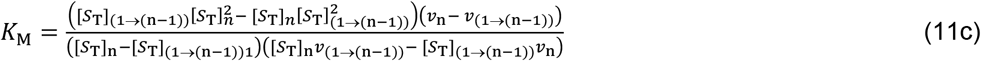

The second possibility is that, if Eq. (9b) is substituted into original Michaelis-Menten equation one gets the equation for *K*_M_ without *V*_max_ given, after simple steps, as:

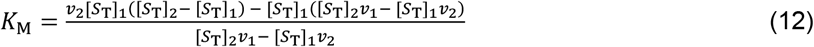

What must not be ignored is that, be it linear regression or nonlinear regression, the curve follows the line of best-fit in order to generate kinetic parameters in which the effect of outliers is minimised on the basis of compromise rather than rectification. Therefore, the parameter generated cannot be seen as being entirely dependent on the original experimental data laden with errors. If there are reasons for the use of the experimental variables obtained from the experiment, then the parameters should be substituted into the original equation, the Michaelis-Menten equation, for instance, in order to calculate those variables, such as velocities corresponding to the measured substrate concentrations, assuming that the measurement was error-free. Alternatively, the substrate concentrations need to be calculated because, peradventure, there may have been inaccurate pipetting of the solution or a mixture of insoluble substrate and solvent. In all these cases, the pipetting of the enzyme solution may be considered error-free. At this juncture, there is a need to point out that something similar, but with a minor difference, in the overall “structure” of the equation is available in the literature, perhaps for the inhibition case [14]. There has always been criticism against any form of regression; surprisingly, nonlinear least squares fitting technique is included [12] despite the application of software. Indeed, software seems unable to correct errors.

## 3.0 MATERIALS AND METHODS

### 3.1 Materials

#### 3.1.1 Chemicals

Aspergillus oryzae alpha-amylase (EC 3.2.1.1) and insoluble potato starch were purchased from Sigma-Aldrich, USA. Tris 3, 5—dinitrosalicylic acid, maltose, and sodium potassium tartrate tetrahydrate were purchased from Kem Light Laboratories in Mumbai, India. Hydrochloric acid, sodium hydroxide, and sodium chloride were purchased from BDH Chemical Ltd., Poole, England. Distilled water was purchased from the local market.

#### 3.1.2 Equipment

An electronic weighing machine was purchased from Wenser Weighing Scale Limited and 721/722 visible spectrophotometer was purchased from Spectrum Instruments, China; pH meter was purchased from Hanna Instruments, Italy.

### 3.2 Methods

#### 3.2.1 Preparation of solution of reactants and assay

The enzyme was assayed according to the Bernfeld method [15] using gelatinised potato starch. Reducing sugar produced upon hydrolysis of the substrate using maltose as a standard was determined at 540 nm with an extinction coefficient equal to 181 L/mol.cm. A concentration equal to 1 g/100 mL of potato starch was gelatinised at 100 °C for 3 min and subjected to serial dilution after making up for the loss of moisture due to evaporation to give concentrations ranging between 4 and 10 g/L for the assay in which [*S*_T_] ≫ [*E*_T_]. A concentration of 0.01 g/100 mL of *Aspergillus oryzae* alpha-amylase was prepared by dissolving 0.01 g of the enzyme (as the stock) in 100 mL of Tris-HCl buffer at pH = 6.9. The assay of the enzyme was carried out with an enzyme concentration of 1 mg/L. The duration of the assay was 3 minutes at 20 °C.

#### 3.2.2 Determination of K_M_ and v_max_ by calculation and graphical method

The determination of *K*_M_ is according to Eqs (4, 13). The ***V***_max_ was obtained by fitting the Eq. (10) to the unweighted velocity data in this experiment and in the literature [4]. Equations (11b), (12), and (9) were left out because of a time constraint; otherwise, the same result is expected using either Eq. (4) or Eq. (10), as the case may be.

The equations (*v*_1_ → *v*_3_) [13] and those derived in this research (*v*_4_ → *v*_9_) used to correct the variables, the velocities (*v*_1_, *v*_2_, *v*_3_, *v*_4_, *v*_5_, *v*_6_, *v*_7_, *v*_8_, and *v*_9_) of enzymatic action, are:

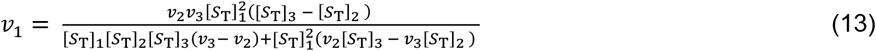

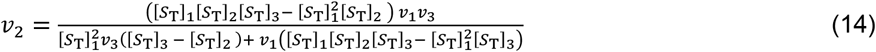

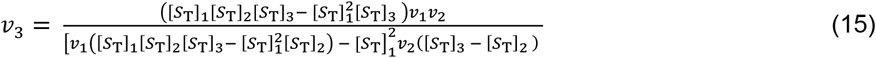

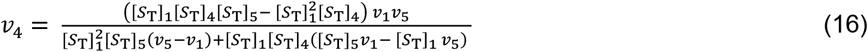

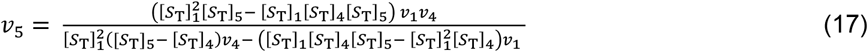

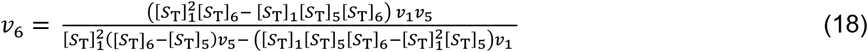

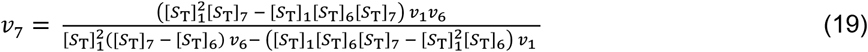

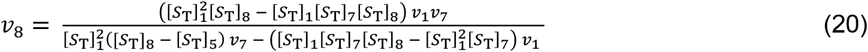

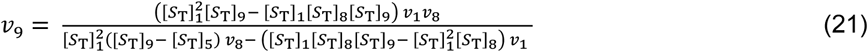

The graphing approaches were a double reciprocal plot and a plot based on Eq. (4) for *K*_M_ and on Eq. (10) for *V*_max_, where respectively, the *x*-axis is taken as *f* ([S], *v*) and the *y*-axis is taken as *f* ([*S*]^2^, *v*), and *f* ([*S*], *v*) and *f* (*v*_2_, [*S*]).

### 3.3 Statistics

Duplicate assays for each substrate were deliberately adopted in this research, not just to reduce time and cost but also to serve as a preliminary test of the mathematical equations derived so as to verify robustness and consistency. As in the previous publication [13], the pseudo-weighting factors for the products and substrates are given in a summarised version as follows:

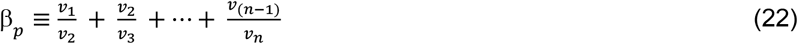

The pseudo-weighting factor for the substrate is given as:

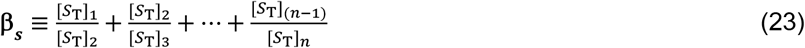

The coefficients, **β**_**s**_ **and β**_**p**_, are taken to be a weighting factor for the fractional contribution of each substrate and each product to the excess (or, generally speaking, the error) observed in the summation results. The summation equations for the *v*_max_ and *K*_M_ are:

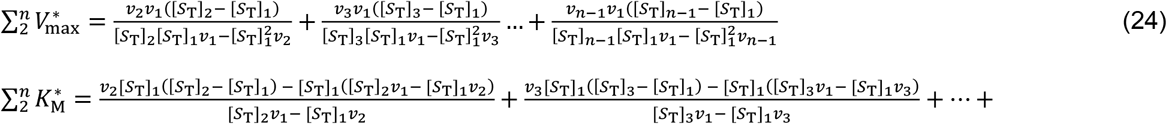

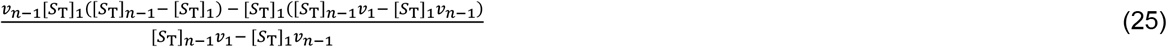

The mathematically and pseudo-statistically determined *V*_max_, *V*_max(p-stat)_, is [13]:

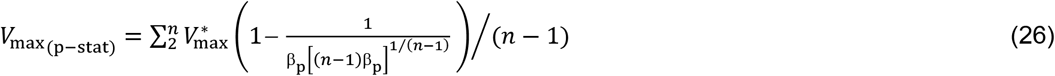

where, *v*_i_ is the original velocity of enzymatic action without weighting or any treatment, and, *n* is the total number of different concentrations of the substrate. The corresponding *K*_M_ is:

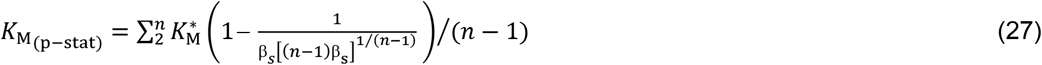

The arithmetic means (AV) are:

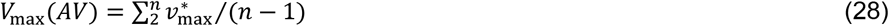

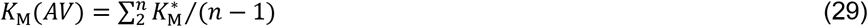

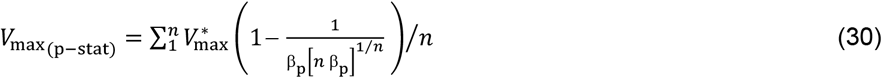

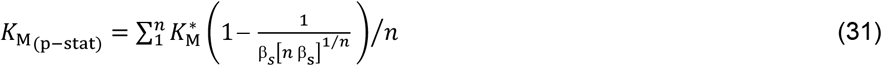

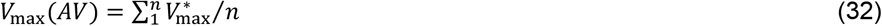

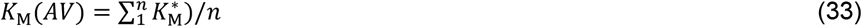

Standard deviations (SD) were calculated using Microsoft Excel with different sample numbers (*n*) for each parameter for different enzymes; values are reported as mean ± SD.

## 4.0 RESULTS AND DISCUUSION

This section is best introduced with an overview of the equations derived in this research. Separate different equations for the calculation of *K*_M_ and *V*_max_ that give the same results are an expression of robustness and consistency, and most importantly, the validity of a procedural issue. To accomplish the goal of validity, the equations had to be evaluated by graphical means, beginning with the double reciprocal plot and then, ten plots based on some of the derived equations (figures 1 → 11). The double reciprocal plots, otherwise called Lineweaver-Burk plots (LWB) [16], using the un-weighted (UNW) and recalculated (RC) initial rates (Table 1), showed that the result in the literature [4], if mistakes are excluded, is far higher than the results shown in figure 1 and Table 2.

**Table 1.** Unweighted and recalculated initial rates or velocities

| **Beta-galactosidase (EC.3.2.1.23)** |  | ***Aspergillus oryzae* alpha amylase (EC. 3.2.1.1)** |  |
| --- | --- | --- | --- |
| UNW velocities data/$\mu\text{M}/\text{min}$ [4] | RCV data/$\mu\text{M}/\text{min}$ | Data given as arithmetic mean of each UNW velocity /$\mu\text{M}/\text{min}$ | RCV data/$\mu\text{M}/\text{min}$ |
| 3 | 3.122449127 | 171.15 | 177.503376 |
| 6 | 6.181818182 | 219.05 | 215.108142 |
| 17 | 15 | 259.55 | 250.4857109 |
| 48 | 27.54545455 | 285.25 | 281.430301 |
| 101 | 90.3919266 | 311.85 | 311.862502 |
| 121 | 126.457077 | 329.45 | 355.606156 |
| 139 | 148.706315 | - | - |
| 152 | 180.4739919 | - | - |
| 181 | 189.4642622 | - | - |

UNW stands for un-weighted velocity. For the benefit of convenience, the substrate, o-nitrophenyl- $\beta$ -D-galactopyranoside (ONPG) concentrations (mM) [4] are: 0.05, 0.1, 0.25, 0.5, 2.5, 5, 8, 20, and 30. Gelatinised insoluble potato starch concentrations (g/L) used for the assay are: 4, 5, 6, 7, 8, and 10. Mean of two determinations was the case for this research as applicable to *A. oryzae* alpha-amylase.

**Table 2.** Michaelian parameters determined according to Lineweaver-Burk method with data in the literature as applicable to EC.3.2.1.23 [4] and in this research (EC 3.2.1.1).

| Beta-galactosidase (EC.3.2.1.23) |  | *Aspergillus oryzae* alpha amylase (EC. 3.2.1.1) |  |
| --- | --- | --- | --- |
| $V_{\max}$ using UNW/$\mu\text{M}/\text{min}$ | 833.333 | $av V_{\max}$ using UNW/$\text{mM}/\text{min}$ | 1.166<br>(1.018) |
| $K_M$ using UNW/$\text{mM}$ | 13.75 | $av K_M$ using UNW/$\text{g/L}$ | 22.323<br>(18.346) |
| $V_{\max}$ using RCV($v_1 \rightarrow v_4$) & UNW($v_5 \rightarrow v_9$)/$\mu\text{M}/\text{min}$ | 243.902<br>(214.097) | $v_{\max}$ using RCV($v_1 \rightarrow v_6$)/$\text{mm}/\text{min}$ | 1.164<br>(1.016) |
| $K_M$ using RCV($v_1 \rightarrow v_4$) & UNW($v_5 \rightarrow v_9$)/$\text{mM}$ | 4<br>(3.311) | $K_M$ using RCV($v_1 \rightarrow v_6$)/$\text{g/L}$ | 22.090<br>(18.154) |
| $V_{\max}$ using RCV ($v_1 \rightarrow v_9$)/$\mu\text{M}/\text{min}$ | 217.391<br>(190.674) | - | - |
| $K_M$ using RCV ($v_1 \rightarrow v_9$)/$\text{mM}$ | 3.435<br>(2.844) | - | - |

UNW and RCV stand for unweighted and recalculated velocities of enzymatic action, respectively. The average of duplicate values of $V_{\max}$ and the average of duplicate values of $K_M$ were taken from two plots. All values in brackets are outcomes of the pseudo-statistical treatment of the kinetic parameters generated from both raw and recalculated or corrected initial rates. The pseudo-statistical factors defined by Eqs (26) and (27) are 0.877791 and 0.8278, respectively, for beta-galactosidase, and 0.872989 and 0.821833, respectively, for alpha-amylase.

**Figure 1:**
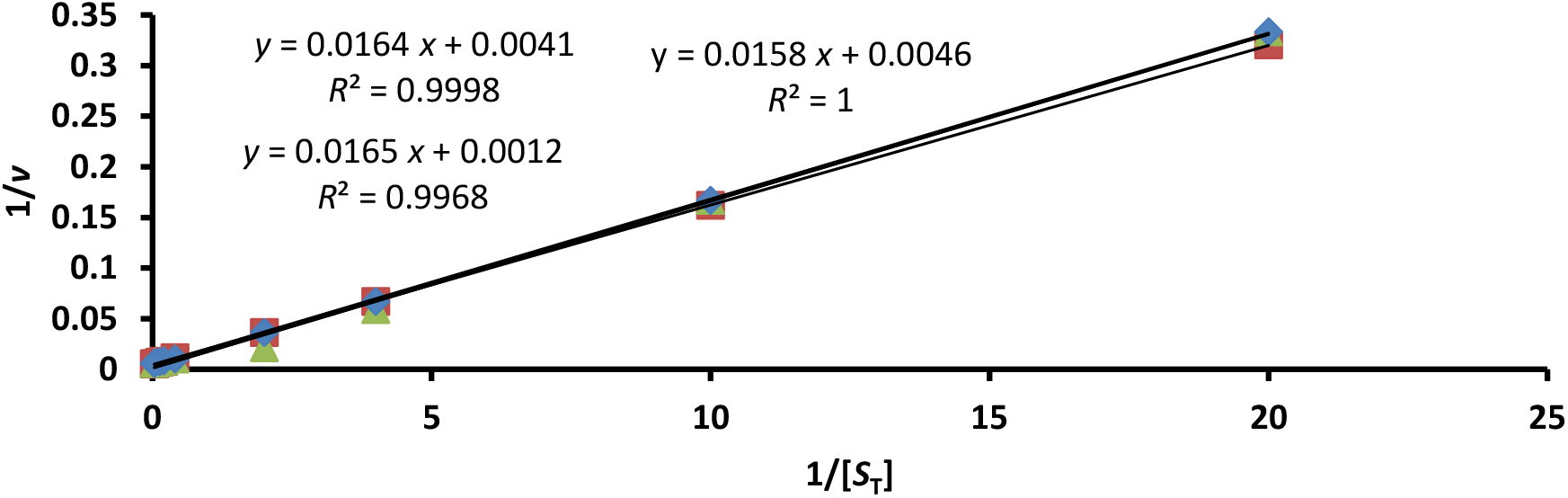
Double reciprocal plot, Lineweaver-Burk plot, using directly original (▴) data (unweighted) in the literature [4] for comparative and confirmation/validation purposes. Other legends are partially recalculated variables 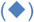, *v*_1_ → *v*_4_, and totally recalculated variables 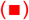, *v*_1_ → *v*_9_. *Relevant equations in this research were fitted to the unweighted data for the purpose of recalculating each velocity, the initial rate, v*_*i*_ *of enzymatic action as may be applicable. The V*_*max*_ *and K*_*M*_ *for the unweighted v*_*i*_ *are respectively 833*.*333 μM/min and 13*.*03 g/L; for the partly corrected vi the values are respectively 243*.*9 μM/min and 4 g/L; for the totally corrected v*_*i*_ *the values are respectively 217*.*391 μM/min and 3*.*435 g/L*.

The figures are deliberately included for immediate visual examination of issues observed or raised; hence, the tables remain complimentary rather than of procedural importance. With the LWB plot, the results (Table 2) were compared as follows: The literature on UNW initial rates [4] with partly RC initial rates plus UNW initial rates (this research) gave kinetic parameters that were **greater than** those given by fully RC initial rates (this research), with correlation coefficients, *R*, ranging between 0.998 and 1 (figure 1). The reported results [4], based on software-assisted nonlinear regression, the *V*_max_ and *K*_M_, were less than what was observed in this research, where values were generated graphically by LWB plots and other plots based on derived equations. The LWB plot for *A. oryzae* was not shown, but the results are shown in Table (2).

Calculated kinetic parameters based on the derived equations are shown in Table 3. It needs to be made clear that those Michaelian parameters (to be emphatic), *K*_M_ and *V*_max_, are functions of total substrate concentration [*S*_T_] and velocity, *v*_i_ of enzymatic action. Hence, the much-discussed transient assays must not only be in terms of time scale; they must also take into account the substrate concentration regime if Michaelian kinetics is in view. If [*S*_T_] range is ≪*K*_M_ and [*S*_T_] ≪[*E*_T_] (total enzyme concentration), the Michaelian formalism (sQSSA) ceases to be relevant, becoming more of a case of rQSSA. In this case, the *v*_*i*_ becomes directly proportional to [*S*_T_]. Under such circumstances, Eqs (4a), (9a), (10), (11b), (12), *etc*. become invalid if intended for the calculation of *K*_M_ or *V*_*max*_, as the case may be. In a perfect direct proportionality, [*S*_T_]_n_*v*_n -1_ − [*S*_T_]_n-1_*v*_n_ = 0, as observed with unweighted data in the literature (see footnote under Table 3); one can even insinuate that [*S*_T_]_n_ *v*_n -1_ − [*S*_T_]_n-1_ *v*_n_ < 0 is better than a zero outcome because at least a negative value of the kinetic parameter would have been achieved, as observed in this research and recorded under Table 3 as a footnote. Both are emphatically invalid. However, such a possibility cannot be ruled out if a single bond substrate is the case, as is applicable to disaccharides and perhaps o-nitrophenyl-b-D-galacto-pyranoside [12]. In general, this may be the case where [*S*_T_] is « [*E*_T_] and [*S*_T_] « *K*_M_ such that *v*_i_ remains directly proportional to [*S*_T_] for at least up to five different [*S*_T_]. In such situation, *v*/[*S*_T_] or [*S*_T_]/*v* for up to five different [*S*_T_] must be constant. But this situation may not be in line with Michaelian kinetics.

**Table 3.**
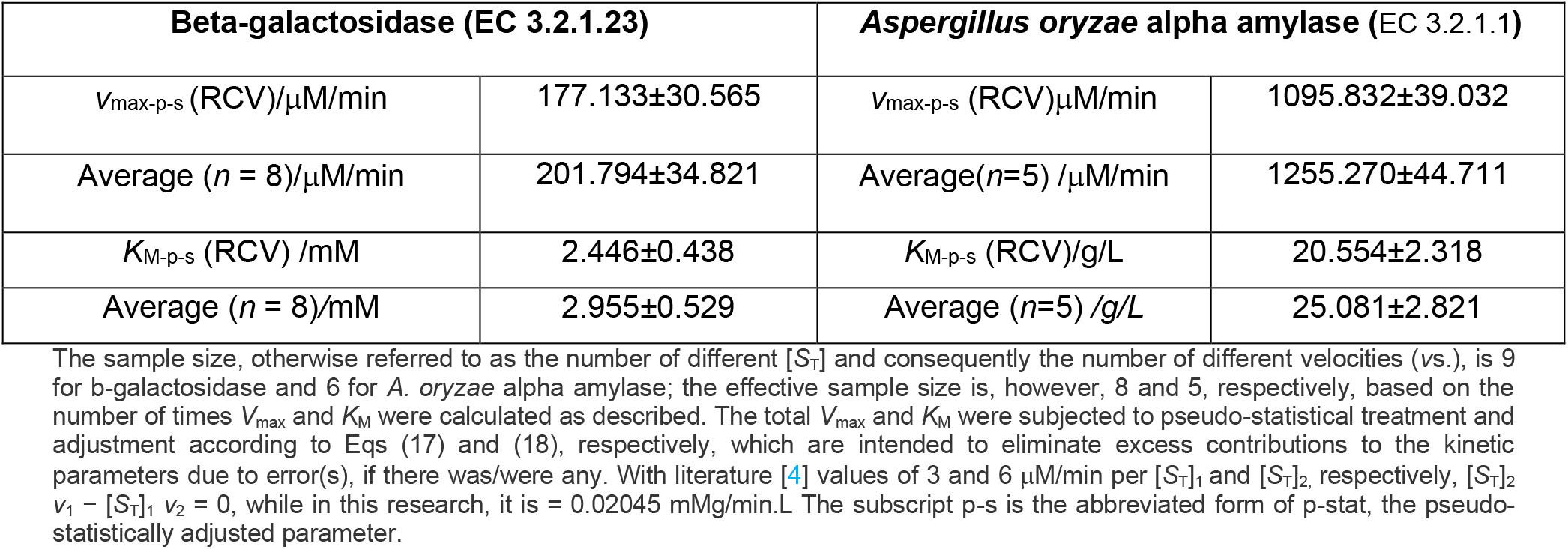
Michaelian parameters, determined by fitting relevant equations in this research to data in the literature (with respect to EC 3.2.1.23 [4]), and in this research (with respect to EC 3.2.1.1).

There is a very strong point in emphasising the need to examine the accuracy of the measured and the experimentally generated variables, *v*_1_, *v*_2_, and *v*_3_, in particular. To achieve this, more specific equations such as Eq. (4a), Eq. (9c), Eq. (12), and Eq. (10), can be used. In all, [*S*_T_]_2_*v*_1_ − [*S*_T_]_1_ *v*_2_ must not yield a negative or zero value. A better value must be greater than 1. The results as quantitative values were obtained first by a double reciprocal plot (figure 1) using data from the literature [4]. Both the raw initial rate data and the corrected version in the literature [4] and in this research are shown in Table 1. All results show that the raw (unweighted) data overestimated kinetic parameters due to the doubling of *v*_i_ with [*S*_T_]_2_, which is also twice the first substrate concentration, [*S*_T_]_1_. As noted elsewhere [5], the other initial rates that did not follow the same pattern were annulled by attenuation rather than total elimination, the effect of the negative intercept. But with partial correction of the initial rates (*v*_1_ → *v*_4_), the kinetic parameters, the *K*_M_ and *V*_*max*_, *are* reduced in magnitude but the values are higher than those for the total corrections, covering the 9 initial rates. The values obtained after the pseudo-statistical treatment (this means multiplying the initial results by a decimal integer defined by Eqs (26) and (27) if necessary) were expectedly lower than the untreated (Table 2). The results, such as 190.7 micro M/min and 2.84 mg/L from the correction of all *v*_i_ values and 214.1 micro M/min and 3.3 mg/L for the partly corrected *v*_i_ values, are not widely different from the literature values of 180 micro M/min and 2.5 mg/L [4].

The values that were overestimated due to the first two initial rates for galactosidase are also due to conditions that invalidate the Michaelis-Menten equation (re-christened in this research as the HBHMM equation) and the associated quasi-steady-state assumption such that [*S*_T_]_n_*v*_n-1_ − [*S*_T_]_n−1_*v*_n_ should be equal to zero. Subjecting such overestimated kinetic parameters as 833.33 *μ*M/min and 13.75 g/L (Table 2) to a pseudo-statistical treatment is ruled out because it is of no value. However, such overestimation cannot be ruled out, assuming accurate values of initial rates, if Michaelian kinetics is out of the question in preference for single turnover kinetics [17]. In general, this may be the case where [*S*_T_] is ≪[*E*_T_] and [*S*_T_] is ≪*K*_M_ such that *v* remains directly proportional to [*S*_T_] for at least up to five different [*S*_T_]. In such a situation, *v*/[*S*_T_] or [*S*_T_]/*v* for up to five different [*S*_T_] must be constant. But this situation may not be in line with Michaelian kinetics.

As a result, it was critical to evaluate the equations, by plotting *f* (*v*^2^, [*S*_T_]) versus *f* (*v*, [*S*_T_]) where the *y*-axis is equivalent to (*v*_n_*v*_n−1_([*S*_T_]_n_ − [*S*_T_]_n-1_)) and *x*-axis is equivalent to ([*S*_T_]_n_*v*_n-1_ − *v*_n_[*S*_T_]_n-1_) and *f* ([*S*_T_]^2^, *v*) versus *f* (*v*, [*S*_T_]) where the *y*-axis is equivalent to ([*S*_T_]_n_[*S*_T_]_n−1_(*v*_n_−*v*_n-1_)) and *x*-axis is equivalent to ([*S*_T_]_n_*v*_n-1_ − *v*_n_[*S*_T_]_n−1_), to yield respectively the *V*_max_ and *K*_M_. All results (equation of linear regression) observed were displayed as an inset and written as a footnote under each corresponding figure, namely, figures 2 and 3 for *A. oryzae* alpha amylase and figures (4 → 7) for beta-galactosidase. The results (1.034 mM/min and 19.296 g/L) from Figures 2 and 3, based on Eqs (9a) and (11b), respectively, were similar to the values (1.016 mM/min and 18.154 g/L) yielded after subjecting the initial results (Table 2) from the LWB plot to a pseudo-statistical treatment. The magnitude of kinetic parameters obtained was < than that obtained by the LWB method.

**Fig. 2:**
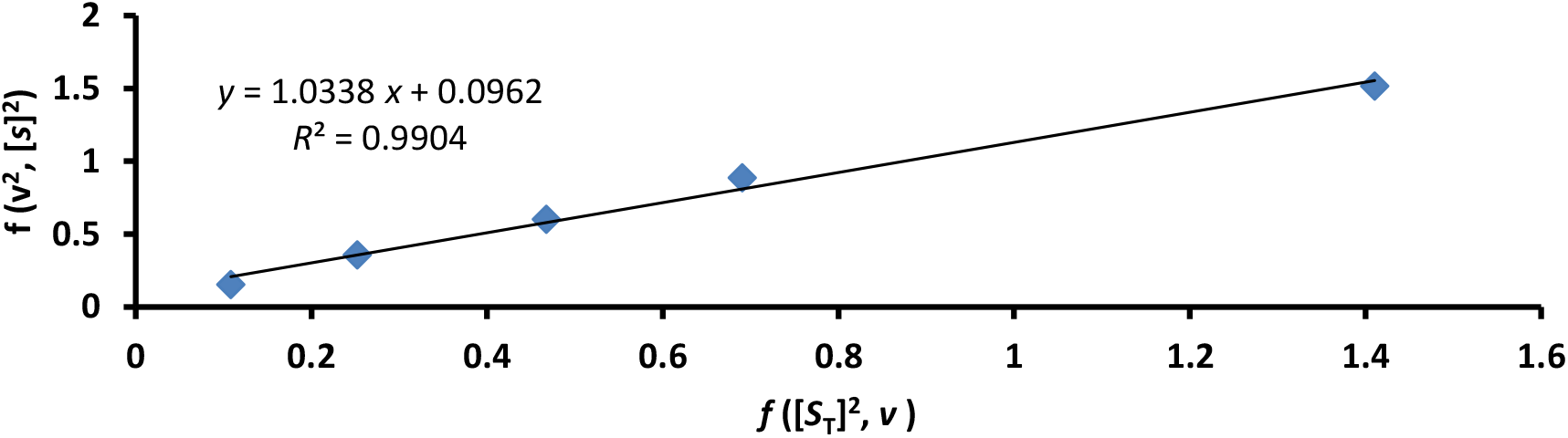
Determination of maximum velocity of enzymatic action, *V*_max_ by graphical method based on Eq. (9a). *The ordinate, y,* 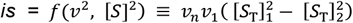 *and the abscissa, x,* 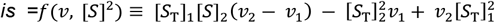: *The inset shows that V*_*max*_ *is = 1*.*034 exp. (*− *3) M/mL/min; R* is ≈ *0*.*99. Data is from this research covering the assay on alpha-amylase*.

**Fig. 3:**
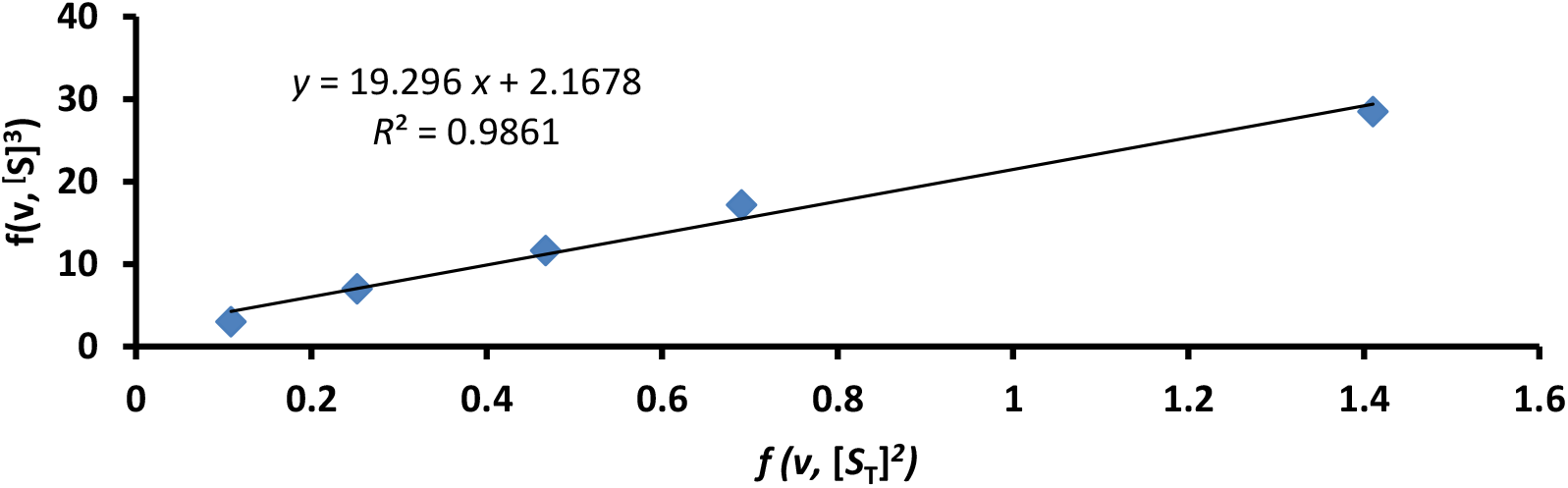
Determination of MM constant, *K*_M_ by graphical method based on Eq. (4/11b) *The ordinate, y,* 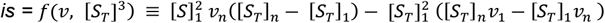 *and the abscissa, x,* 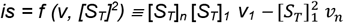: *The inset shows that K*_*M*_ *is = 19*.*296 g/L; R is* ≈ *0*.*99. Data is from this research*.

The second set of plots (figures (8) → (11)) were plots of *v*_n_*v*_i_ ([*S*_T_]_n_ − [*S*_T_]_i_) versus [*S*_T_]_n_ *v*_i_ − [*S*_T_]_i_*v*_n._ Fitting the equations to the recalculated variables (the velocities) and then plotting gave magnitudes of values that were < those observed in LWB plots, with *R* values being perfectly = 1 in one instance. However, such values were not widely different from those obtained from weighted linear and nonlinear regression in the literature [4]. The results garnered from using partly corrected *v*_i_ values (*v*_1_→*v*_4_) and fully corrected *v*_i_ values are quite lower than the values garnered from a plot (LWB plot) using the raw data. The values of the parameters *V*_max_ (figure 4 and Eq. (9c)) and *K*_M_ (figure 5 and Eq. (4a) as percentages of inaccurate parameters are respectively 23.08 and 22.03 %; the pseudo-statistically remediated values are 168.83 mM/min and 2.508 g/L, which correspond to the initial measurements of 192.33 *μ*M/min and 3.03 g/L, respectively. Here, one sees that the initial measurements were not overestimates, even if they were > than those in the literature report. The literature report [4], however, reveals a burden of error in the initial rates, which may have been attenuated by the mechanism and assumptions of the nonlinear regression software package. One must, however, admit that only one of the eight replicates of the initial rates was made available in the literature. It was sufficiently useful for the illustration of the facts and principles advanced in this research.

**Figure 4:**
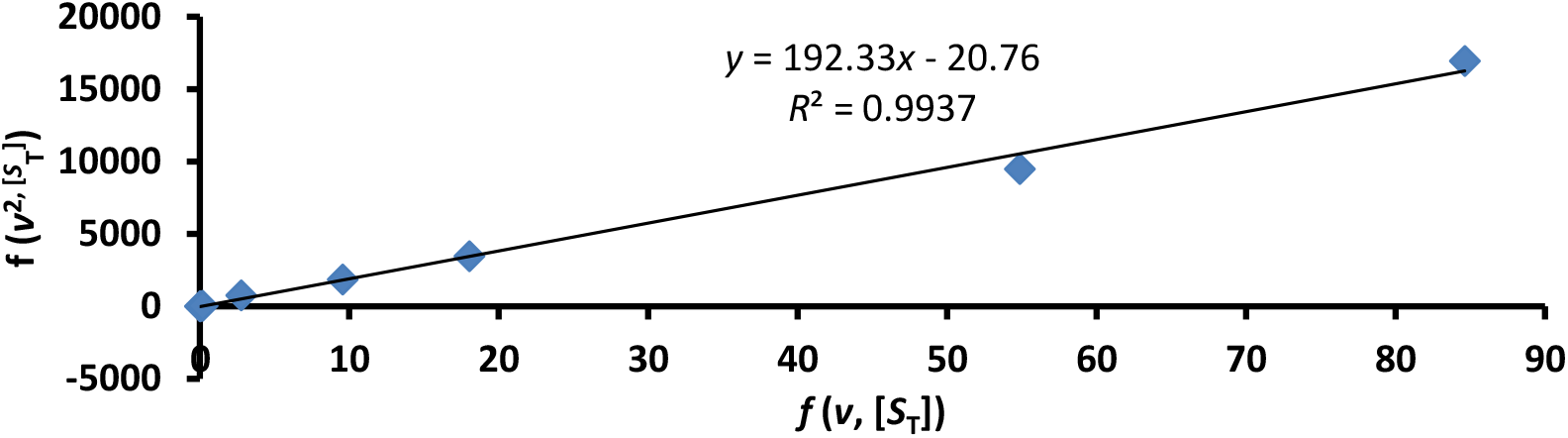
Determination of maximum velocity of enzymatic action, *V*_max_ by graphical method based on Eq. (9c). *The ordinate, y*, is *=f*(*v*^2^, [*S*]) ≡ *v*_*n*_*v*_*n*−1_([*S*]_*n*_ − [*S*]_*n*−1_) *and the abscissa, x, is =f*(*v*, [*S*]) ≡ [*S*]_*n*_*v*_*n*−1_ − [*S*]_*n*−1_*v*_*n*_: *The inset shows that V*_*max*_ *is* ≈ *192*.*33 μM/min (23*.*08 % of the inaccurate value); R is* ≈ *0*.*99. The pseudo-statistically remediated value is 168*.*826 μM/min. The original velocities, v*_*1*_, *v*_*2*_, *v*_*3*_, *and v*_*4*_ *were recalculated according to corresponding equations, Eq. (13)* → *Eq. (16). The original data explored is in the literature [4]*.

**Fig. 5:**
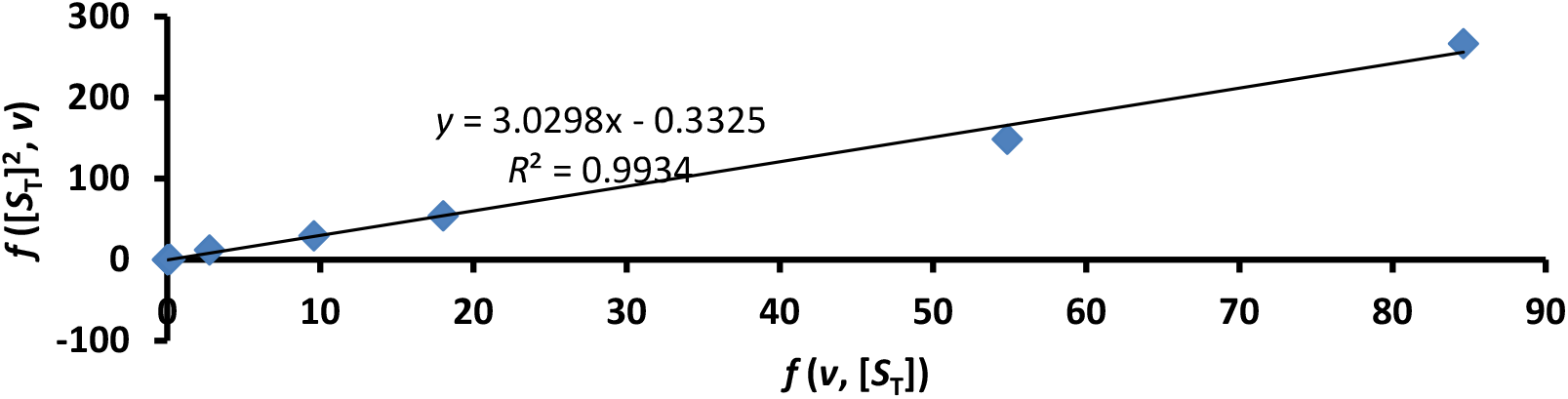
Determination of MM constant, *K*_M_ by graphical method based on Eq. (4a). *The ordinate, y, is = f*(*v*, [*S*]^2^) ≡ [*S*_T_]_*n*_[*S*_T_]_*n*−1_(*v*_*n*_ − *v*_*n*−1_) *and the abscissa, x, is = f (v, [*S_T_]) ≡ *[S*_*T*_]_*n*_ *v*_*n−1*_ − *[S]*_*1*_*v*_*n−1*_ *the inset shows that K*_*M*_ *is* ≈ *3*.*03 mM (22*.*04 % of inaccurate value); R is* ≈ *0*.*99. The pseudo-statistically remediated value is 2*.*508g/L. The data explored is in the literature. The original velocities, v*_*1*_, *v*_*2*_, *v*_*3*_, *and v*_*4*_ *[4] were calculated according to corresponding equations, Eq. (13)* → *Eq. (16)*.

Using all corrected *v*_i_ values, the values of the parameters garnered, *K*_M_ (figure 6 and Eq. (4a)) and *V*_max_ (figure 7 and Eq. (9c) as percentages of inaccurate parameters, are respectively 24.35 and 22.35 %; the pseudo-statistically remediated values are 163.52 micro M/min and 2.508 g/L, which correspond to the initial measurement of 186.285 micro M/min and 3.348 g/L, respectively. Here, one sees that the initial measurements were not overestimates, even if they were greater than those in the literature report. Using figure 8 and Eq. (9d) for *V*_max_ and figure 9 and Eq. (4b) for *K*_M_, coupled with the use of all corrected initial rates, the values of the parameters as percentages of inaccurate parameters are, respectively, 25.28 and 24.20 %; the pseudo-statistically remediated values are 184.687 micro M/min and 2.755 g/L, which correspond to the initial measurements of 3.348 g/L and 186.285 micro M/min.

**Figure 6:**
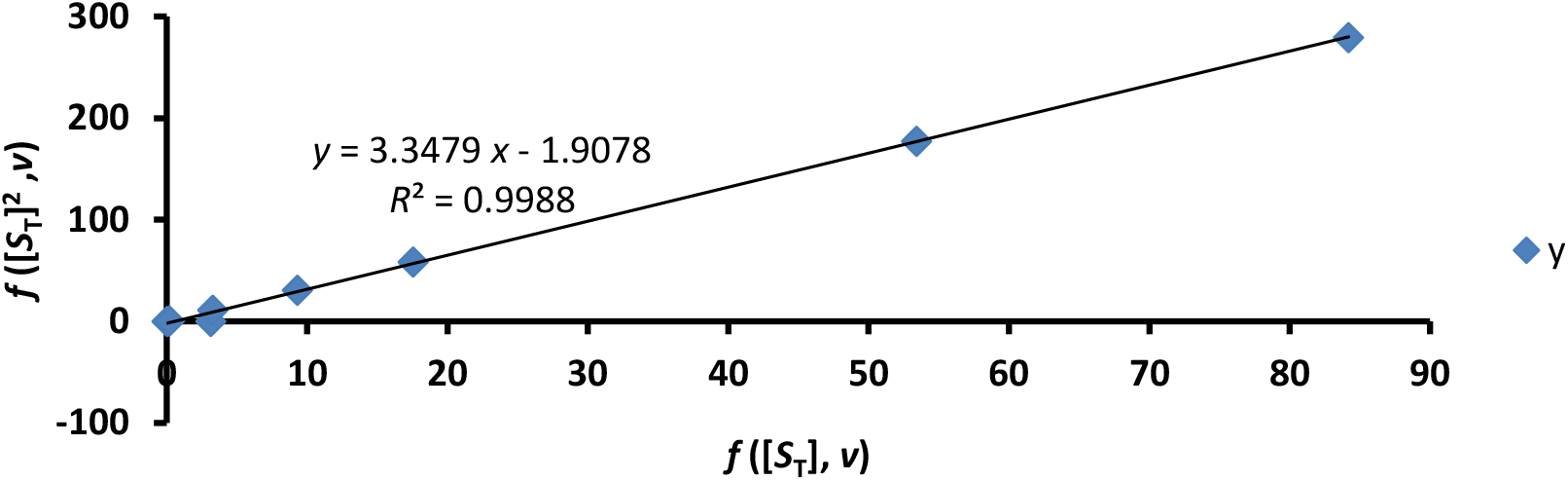
Determination of MM constant, *K*_M_ by graphical method based on Eq. (4a). *The ordinate, y, is = f*(*v*, [*S*_T_]^2^ ≡ [*S*_T_]_*n*_[*S*_T_]_1_(*v*_*n*_ − *v*_1_) *and the abscissa, x, is = f (v, [S*_T_]) ≡ *[S*_T_]_*n*_ *v*_*1*_ − *[S*_T_]_*1*_*v*_*n*_: *The inset shows that K*_*M*_ *is* ≈ *3*.*348 mM (22*.*04 % of the inaccurate value); R is* ≈ *0*.*999. The data explored is in the literature. The pseudo-statistically remediated value is 2*.*772 g/L. The original velocities, v*_*1*_, *v*_*2*_, *v*_*3*_, *v*_*4*_, *v*_*5*_, *v*_*6*_, *v*_*7*_, *v*_*8*_, *and v*_*9*_ *[4] were recalculated according to corresponding equations, Eq. (13)* → *Eq. (21)*.

**Fig. 7:**
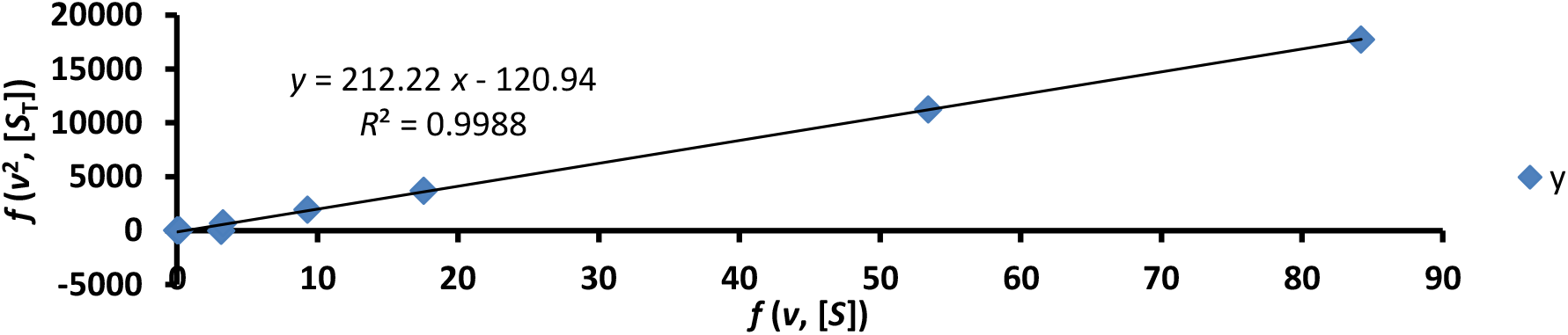
Determination of maximum velocity of enzymatic action, *V*_max_ by graphical method based on Eq. (9c). *The ordinate, y, is = f*(*v*^2^, [*S*_T_]) ≡ *v*_*n*_*v*_1_([*S*_T_]_*n*_ − [*S*_T_]_1_) *and the abscissa, x, is =f*(*v*, [*S*_T_]) ≡ [*S*_T_]_*n*_*v*_1_ − [*S*_T_]_1_*v*_*n*_: *The inset shows that V*_*max*_ *is* ≈ *212*.*22 μM/min* (22.354 % of the inaccurate value); *R is* ≈ 0.999. *The pseudo-statistically remediated value is 163*.*519 μM/min. The original velocities, v*_*1*_, *v*_*2*_, *v*_*3*_, *v*_*4*_, *v*_*5*_, *v*_*6*_, *v*_*7*_, *v*_*8*_, *and v*_*9*_ *were recalculated according to corresponding equations, Eq. (13)* → *Eq. (21). The original data explored is in the literature [4]*.

**Fig. 8:**
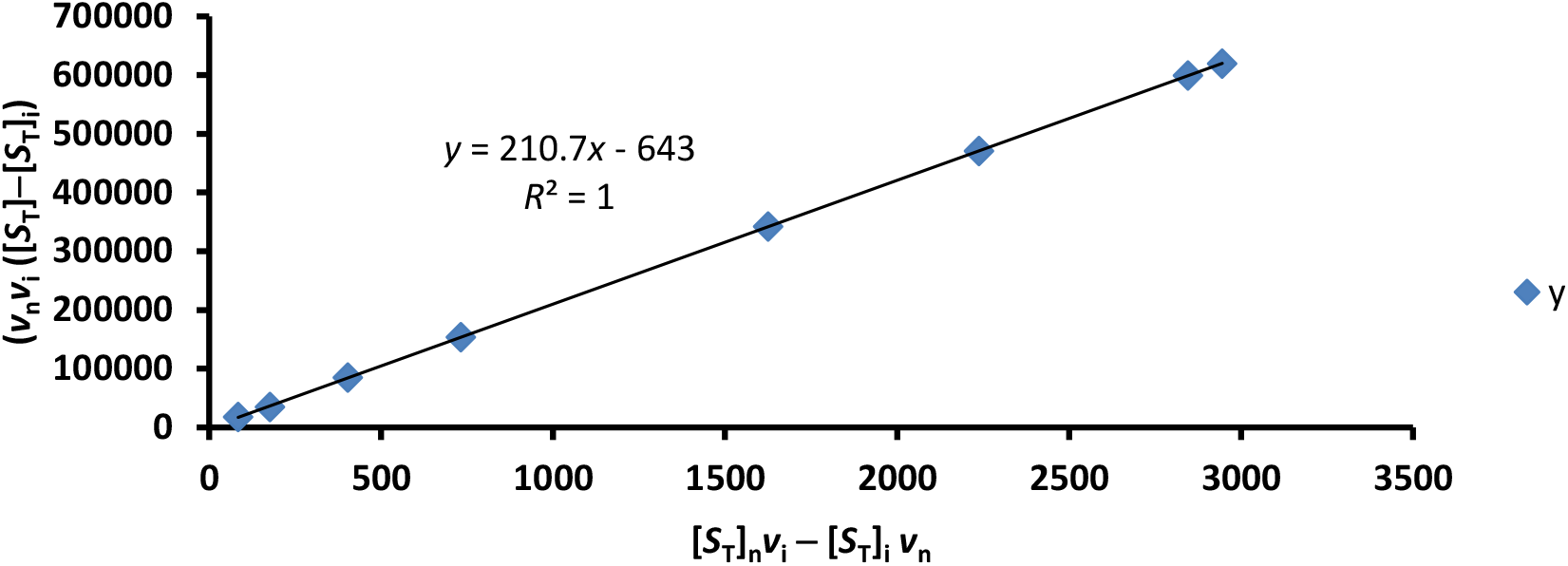
Determination of maximum velocity of enzymatic action, *V*_max_ by graphical method based on Eq. (9d). *The ordinate, y, is = f*(*v*^2^, [*S*_*T*_]) ≡ *v*_*n*_*v*_*i*_([*S*_*T*_]_*n*_ − [*S*_*T*_]_*i*_) *and the abscissa, x, is = f*(*v*, [*S*_*T*_]) ≡ [*S*_*T*_]_*n*_*v*_*i*_ − [*S*_*T*_]_*i*_*v*_*n*_; *i is always = 1. The inset shows that V*_*max*_ *is* ≈ *210*.*7 μM/min (25*.*284 % of the inaccurate value); R is = 1. The pseudo-statistically remediated value is 184*.*687 μM/min. The original velocities, v*_*1*_, *v*_*2*_, *v*_*3*_, *v*_*4*_, *v*_*5*_, *v*_*6*_, *v*_*7*_, *v*_*8*_, *and v*_*9*_ *were recalculated according to corresponding equations, Eq. (13)* → *Eq. (21). The original data explored is in the literature [4]*.

**Fig. 9:**
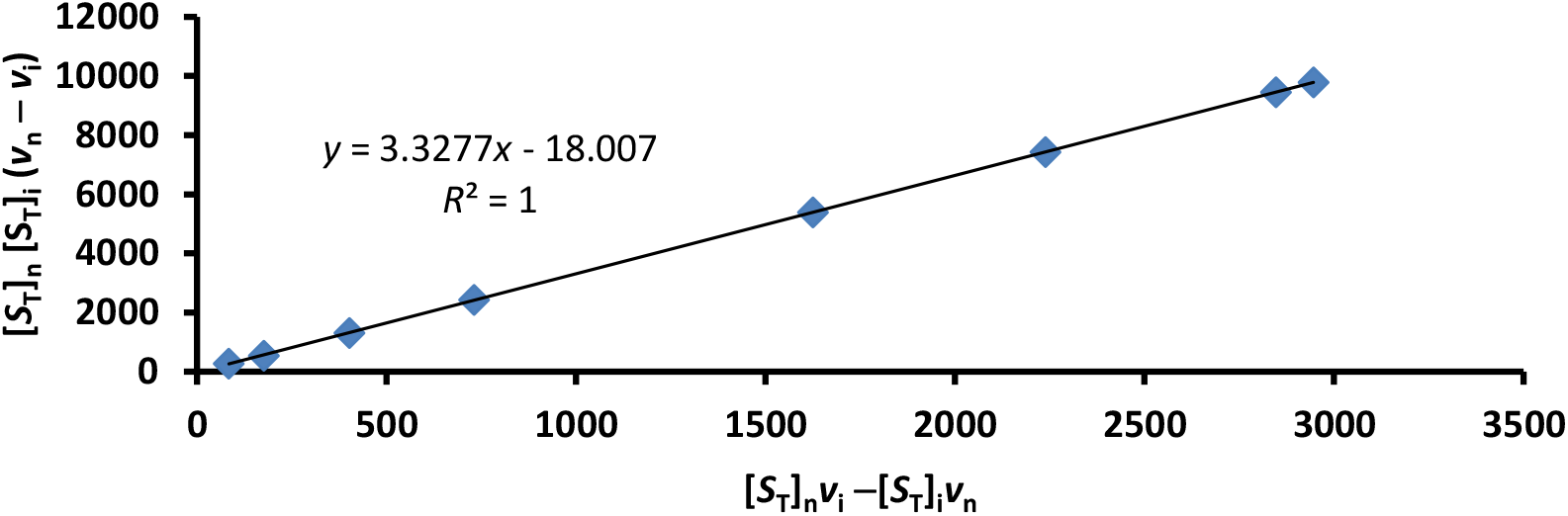
Determination of MM constant, *K*_M_ by graphical method based on Eq. (4b). *The ordinate, y, is = f*(*v*, [*S*]^2^) ≡ [*S*_T_]_*n*_[*S*_T_]_i_(*vn* − *v*i) *and the abscissa, x, is = f (v, [S*_*T*_]) ≡ *[S*_T_]_*n*_ *v*_*i*_ − *[S*_T_]_*i*_*v*_*n*_: *The inset shows K*_*M*_ ≈ *3*.*3277 mM (23*.*528 % of inaccurate value); R is = 1. The pseudo-statistically remediated value is 2*.*755 g/L. The data explored is in the literature. The original velocities [4], v*_*1*_, *v*_*2*_, *v*_*3*_, *v*_*4*_, *v*_*5*_, *v*_*6*_, *v*_*7*_, *v*_*8*_, *and v*_*9*_ *were recalculated according to corresponding equations, Eq. (13)* → *Eq. (21)*.

Timing error does not just arise because of failure to terminate reactions consistently; it also arises if the duration of the assay is such that it totally depletes the substrate before the expiry of the time regime where the lower end of the concentration is the case, but if the upper range of the concentration is the case, the reaction continues until termination by the experimenter. This amounts to a timing error. It does not matter if the duration is on the millisecond time scale. The equations given in this research serve to correct such errors in kinetic variables for the first three assays at three different concentrations of the substrate, as noted in the literature [13]. It is not certain whether computer software can make such adjustments or corrections. Besides, the question is (though in a different context): “Is there anything left to say on enzyme kinetic constants and quasi-steady state approximation?” [18], seems to be given a partial answer in this research. There may be more to say yet.

Surprisingly, fitting the equations to the unweighted data and plotting the results yielded values (2.498 mM and 196.07 mM/min) that are greater than those obtained using the recalculated velocity data, but with an abysmally low correlation coefficient, *R* (0.474) with respect to *K*_M_. The *K*_M_ was, therefore, similar to the 2.5 mM obtained by weighted linear and nonlinear regression in the literature. This is a pointer to the efficacy of the equations. It must be emphasised again that the values do not represent the ultimate high precision value, but rather a substantial improvement. Thus, using Figure 10 and Eq. (9d) for *V*_max_ and figure 11 and Eq. (4b) for *K*_M_, coupled with the use of all unweighted initial rates, the values of the parameters as percentages of inaccurate parameters are, respectively, 23.528 and 18.16 %; the pseudo-statistically remediated values are 172.108 mM/min and 2.067 g/L, which correspond to the initial measurements of 196.07 mM/min and 2.498 g/L, respectively.

**Figure 10:**
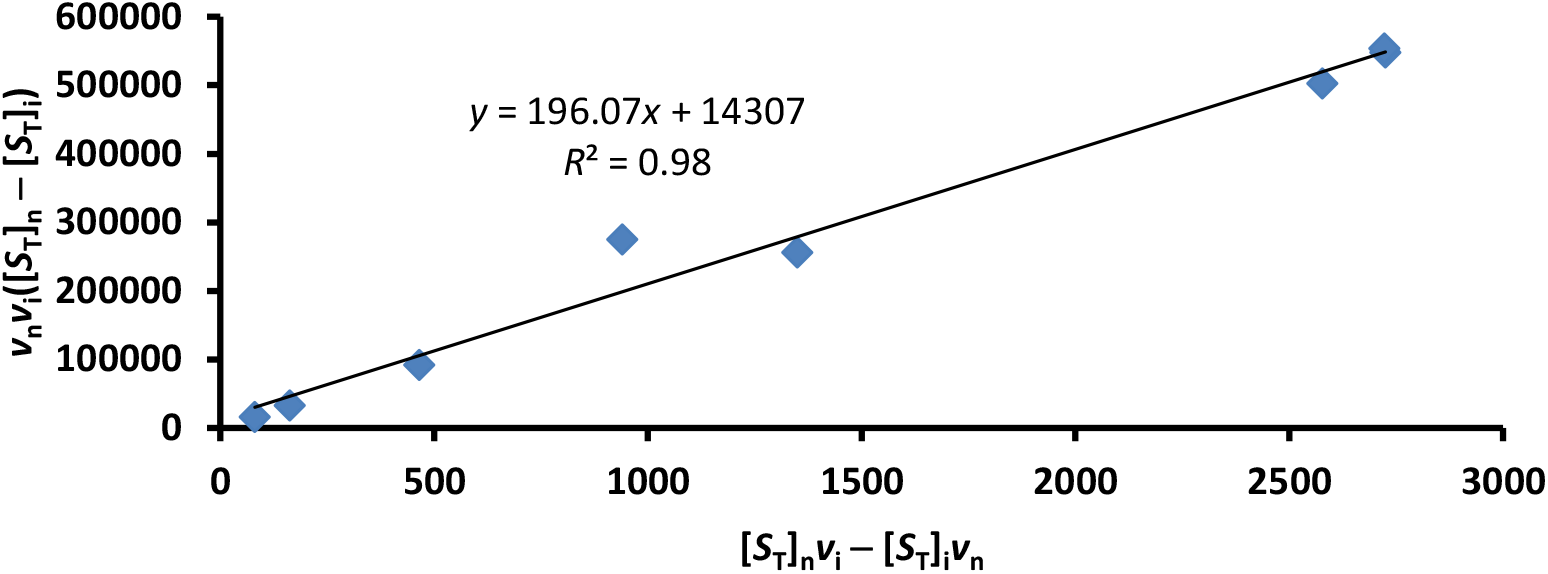
Determination of maximum velocity of enzymatic action, *V*_max_ by graphical method based on Eq. (9d). *The ordinate, y, is = f*(*v*^2^, [*S*_T_]) ≡ *v*_*n*_*v*_*i*_([*S*_T_]_*n*_ − [*S*_T_]_*i*_) *and the abscissa, x, is =f*(*v*, [*S*_T_]) ≡ [*S*_T_]_*n*_*v*_*i*_ − [*S*_T_]_*i*_*v*_*n*_; *i is always* = *1. The inset shows that V*_*max*_ *is = 196*.*07 μM/min (23*.*528 % of the inaccurate value); R is* = *1. The pseudo-statistically remediated value is 172*.*108 μ*M/min. *The original velocities (unweighted), v*_*1*_, *v*_*2*_, *v*_*3*_, *v*_*4*_, *v*_*5*_, *v*_*6*_, *v*_*7*_, *v*_*8*_, *and v*_*9*_ *were used. The original data explored is in the literature [4]*.

**Fig. 11:**
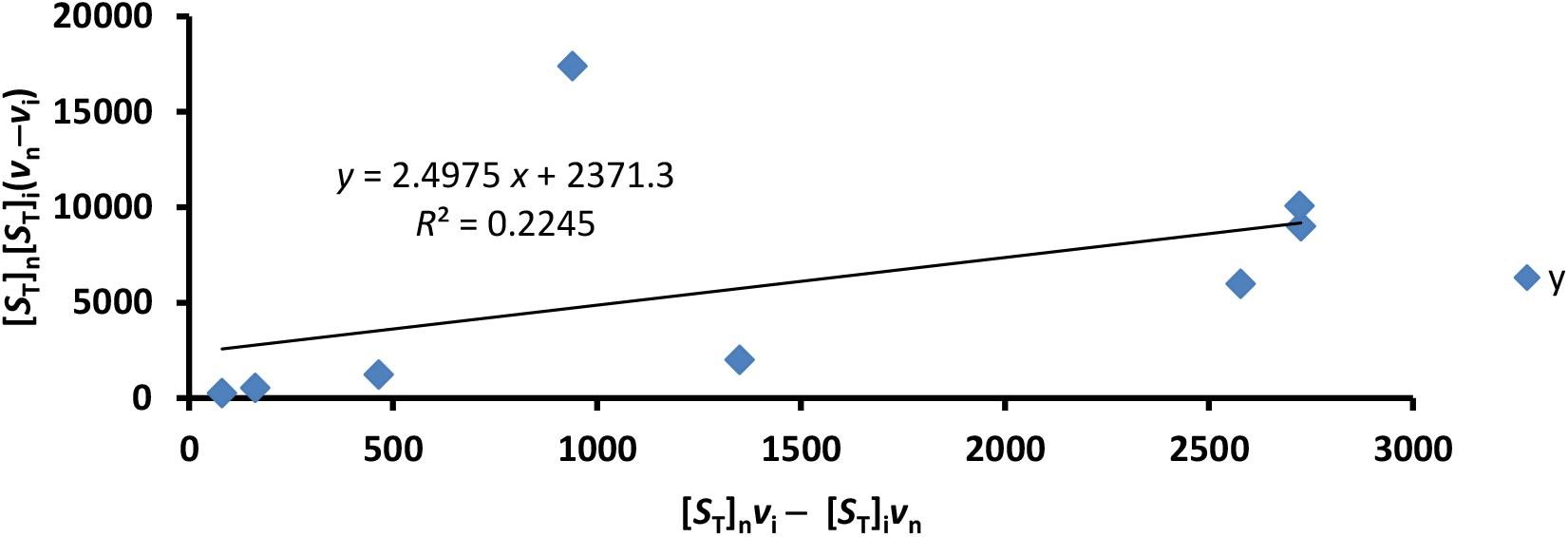
Determination of MM constant, *K*_M_ by graphical method based on Eq. (4b). *The ordinate, y, is = f*(*v*, [*S*_T_]^2^) ≡ [*S*_T_]_*n*_[*S*_T_]_*i*_(*v*_*n*_ − *v*_*i*_) *and the abscissa, x, is = f (v, [S*_T_]) ≡ *[S*_T_]_*n*_ *v*_*i*_ − *[S*_T_]_*i*_*v*_*n*_: *The inset shows that K*_*M*_ *is* ≈ *2*.*498 mM (18*.*16% of inaccurate value); R is* ≈ *0*.*474. The pseudo-statistically remediated value is 2*.*067 g/L. The data explored is in the literature. The original velocities (un-weighted), v*_*1*_, *v*_*2*_, *v*_*3*_, *v*_*4*_, *v*_*5*_, *v*_*6*_, *v*_*7*_, *v*_*8*_, *and v*_*9*_ *[4] were used*.

The outcome of this study notwithstanding, one must bear in mind that if there is no error in all measurements (be it 8 or more replicates for each substrate) under conditions that justify the Michaelian equation and underlying assumptions, there cannot be any need for statistical remediation for generating kinetic parameters; thus, the requirement for statistical soundness and absence of any calculation is out of the question [9]. As opined in a recent preprint report [19], there may be calculations depending on the approach to the solution to any problem of interest. For instance, what has been regarded as the best form of the kinetic parameter, the specificity constant (SC), must be calculated given a single intersection in a reciprocal variant of the direct linear plot by taking the reciprocal of the ratio of *K*_M_ to *V*_max_. But if errors are inevitable even with the use of high-tech devices, then the initial rates must be subjected to correctional treatment, which should ultimately reduce the number of intersections to a minimum.

## 5. CONCLUSION

The equations for the determination of the *K*_M_ and *V*_max_, which are respectively invariant with respect to each other, were rederived. These were in addition to other equations for the same purpose and for the correction of initial rates. The recalculated (or corrected) initial rates gave results for kinetic parameters by graphical means, the LWB method, linear regression based on derived equations, and calculations based on derived equations, which represent a remarkable improvement on the LWB- generated results using unweighted results. The *V*_max_ and *K*_M_ values for galactosidase by graphical means respectively range between 163 and 185 mM/min and between 2.07 and 2.77 g/L; the ranges by calculations are 177 and 214 mM/min and 2.45 and 3.311 g/L, subject to pseudo-statistical remediation.

Overall, the ranges of *V*_max_ and *K*_M_ values for alpha-amylase from both the graphical method and calculation are, respectively, 1.095 to 1.018 mM/min and 18.15 to 20.554 g/L. Nonetheless, the underlying issue remains the conditions that validate Michaelian or non-Michaelian kinetics for the generation of kinetic parameters. The initial rates must not be a mixture of both if the true *K*_M_ and *V*_max_ are of interest. The new pseudo-statistical method for the remediation of error in all measurements, if necessary, is viable, useful, and robust. A future study should examine the effect of high-precision instrumentation for assay in conditions that validate specified QSSA so as to verify the desirability of any statistical approach for the remediation initial rates and kinetic parameters in particular.

## AUTHOR CONTRIBUTION

The sole author designed, analysed, interpreted and prepared the manuscript.

## ACKNOWLEDGMENT

The management of the Royal Court Yard Hotel in Agbor, Delta State, Nigeria, is immensely appreciated for the supply of electricity during the preparation of the manuscript. The provider of the QuillBot grammar checker is thanked for improving the English language quality of the manuscript.

## FUNDING

Funding was privately provided.

## COMPETING INTEREST

There is no competing interest; no financial interest with any government or corporate body or any individual except the awaited unpaid retirement benefits by the Delta state government of Nigeria, for about four years after retirement.

## Notes

### Competing Interest Statement

The authors have declared no competing interest.

